# Depth-Sensitive Assessment of Cerebral Blood Flow and Low-Frequency Oscillations After Traumatic Brain Injury in Mice Using Time-Gated Diffuse Correlation Spectroscopy

**DOI:** 10.1101/2025.05.16.654521

**Authors:** Sahar Sabaghian, Chien-Sing Poon, Dharminder S. Langri, Timothy M. Rambo, Aaron J. Miller, Brandon Foreman, Ulas Sunar

**Affiliations:** Dept of Biomedical Engineering, Stony Brook University, Stony Brook, NY, USA; Dept of Biomedical Engineering, Wright State University, Dayton, OH, USA; Quantum Opus LLC; Dept of Neurology & Rehabilitation Medicine, University of Cincinnati, Cincinnati, OH, USA

**Keywords:** cerebral blood flow, traumatic brain injury, time-gated diffuse correlation spectroscopy (TG-DCS), early and late gate, optical blood flow

## Abstract

Traumatic brain injury (TBI) can lead to long-lasting impairments in cerebral perfusion, making early detection of microvascular changes critical for guiding clinical interventions. In this study, we employed time-gated diffuse correlation spectroscopy (TG-DCS) at 1064 nm to non-invasively quantify depth-resolved cerebral blood flow (CBF) and low-frequency oscillations (LFOs) in a mouse model of closed-head injury. By analyzing early (superficial) and late (deeper) photon arrival times, we identified a significant drop in CBF shortly after injury, with a partial recovery observed at 2 hours post-trauma. Power spectral analysis of the blood flow index revealed significant alterations in LFO bands, particularly in slow-5 (0.01–0.027 Hz) and slow-3 (0.073-0.198 Hz) ranges, with p < 0.05 at both early and late gates. These changes were more pronounced than BFI alterations alone, indicating that LFOs may serve as sensitive biomarkers of neurovascular disruption. Our findings demonstrate the feasibility of TG-DCS for depth-specific monitoring of cerebral hemodynamics and oscillatory dynamics after TBI and suggest its potential utility in translational neurotrauma research.

## 1. Introduction

Traumatic brain injury (TBI) is a major global health concern and a leading cause of death and disability, particularly in young adults ^1^. While primary injury occurs at the time of trauma, secondary injury processes, such as impaired autoregulation, ischemia, and neuroinflammation, can evolve over minutes to hours and significantly affect outcomes ^2^ caused or exacerbated by traumatic blood flow dysregulation ^3^. Early detection of these secondary changes is essential for timely intervention and improved prognosis ^4^. Thus, continuous monitoring of cerebral blood flow (CBF) is an important measure that can be used to detect and prevent these secondary injuries.

Traditional imaging techniques such as MRI, CT, and PET offer detailed measurements of CBF, but are costly, time-consuming, and not well-suited for continuous bedside monitoring. Diffuse correlation spectroscopy (DCS) offers a promising non-invasive optical approach to monitor CBF at the bedside. DCS is a recent optical technique that provides non-invasive, continuous monitoring of CBF, especially useful for high-risk populations. DCS has shown promise in distinguishing cerebral from scalp blood flow, making it valuable TBI monitoring. Compared to NIRS, DCS has higher brain sensitivity ^5–12^. Traditional continuous wave (CW) DCS systems face limitations in measuring absolute blood flow and can be prone to contamination coming from superficial signals such as the skull and scalp. Recent advancements like time-domain DCS (TD-DCS)^13–15^ address these issues by enabling deeper tissue probing and higher signal-to-noise ratios using longer wavelengths, such as 1064 nm, mainly due to lower tissue scattering and higher permissible laser power ^16–21^.

To investigate the early changes in a depth-sensitive manner, we applied the time-gating (TG)-DCS in a mouse model of traumatic brain injury (TBI) induced by a weight-drop. This well-established model allows the study of neurophysiological deficits and cell death associated with TBI *in vivo*. The first objective of this research is to quantify the early alterations in blood flow using time-gated DCS non-invasively and frequently. Growing evidence suggests that low-frequency oscillations (LFOs) in cerebral hemodynamics, typically below 0.2 Hz, may reflect the functional status of neurovascular coupling and autoregulation ^22–46^. Alterations in LFO power spectra have been linked to TBI, stroke, and other neurovascular pathologies, yet their depth-specific features remain underexplored. Thus, as our second objective, we analyzed the power spectrum density of low-frequency oscillations (LFOs) that occur within the blood flow signal and have been linked to human and animal vascular health ^30,44,45^. TBI disrupts cerebral autoregulation (CA), damages neurovascular structures, and induce changes in functional connectivity networks that are linked by these low-frequency oscillations (LFOs) ^31,47,48^. We investigated the capability of time-gated diffuse correlation spectroscopy (TG-DCS) for detecting depth-sensitive changes in these spontaneous LFOs arising from TBI as potential biomarkers of traumatic neurovascular injury.

## 2. Results

### 2.1. Time-domain characteristics

The first objective of this study was to assess the early changes in BFI obtained from three time points (T0, T30, and T120). We first examined the time-series data of blood flow index (BFI) across all TBI mice in the early and late gate. **Figure 2** shows the representative results measured by TG-DCS. **Figures 2(a)** show the temporal point spread function (TPSF) curves from a single measurement with the early and late gates highlighted. The early gate opens at -100ps with a width of 20ps and the late gate opens at +350ps with a width of 200ps from the peak (red overlay). **Figure 2(b)** demonstrates the g2 curves of representative data that are fitted with the time-gated g2 model in both the early and late gates, respectively. **Figure 2(c)** represents the measurement of mean BFI changes pooled across all animals in sham (control) and injury groups at pre-injury (T0) and post-30 minutes (T30) and post-120 minutes (T120) for both gates. For the early-gate, the sham group had baseline (T0): ∼0.065±0.02, T30: ∼0.067±0.02 and T120: ∼0.066±0.02, and for the late-gate, T0: ∼0.20±0.03, T30: ∼0.20±0.03, and T120: ∼0.19±0.03.

**Figure 1.**
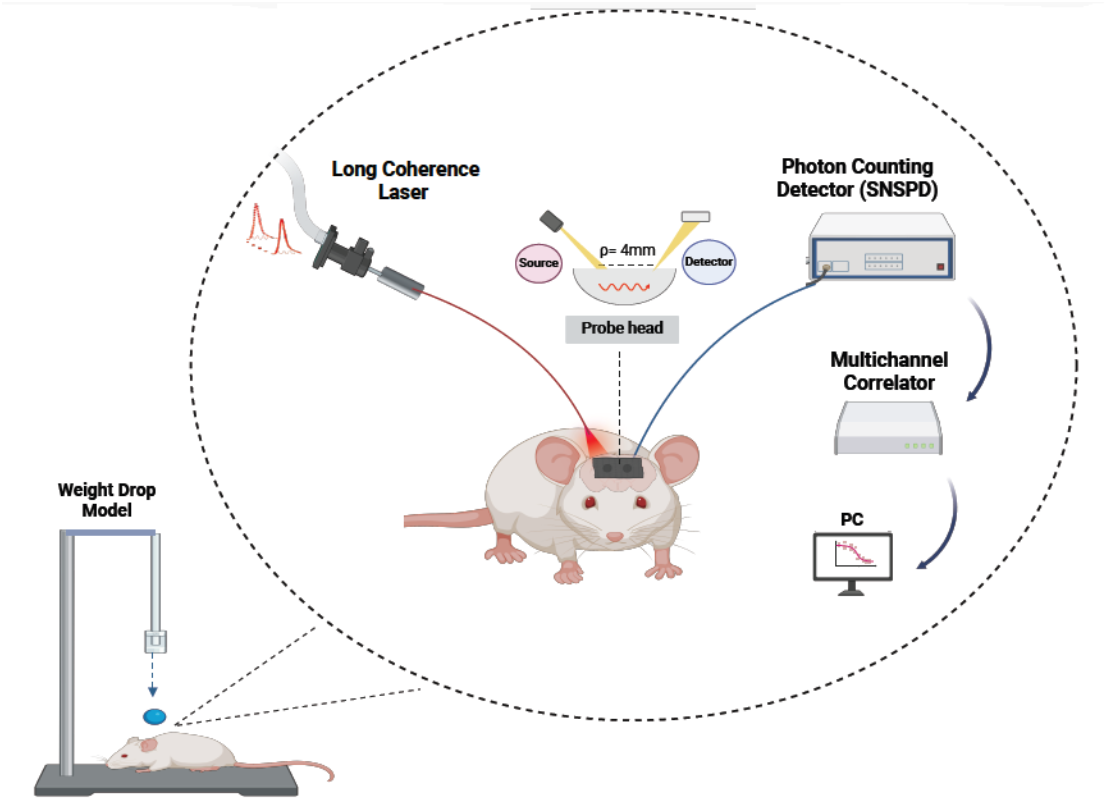
The diagram of the weight-drop and experimental setup. A 1064nm coherent laser emitted and detected by an SNSPD detector through fibers on the probe positioned on the scalp. The source-detector (s-d) separation was 4 mm. The temporal point spread function (TPSF) is used for time-gating to separate superficial vs. deeper signal, and the software autocorrelation is calculated for each gate to assess blood flow index (BFI).

**Figure 2.**
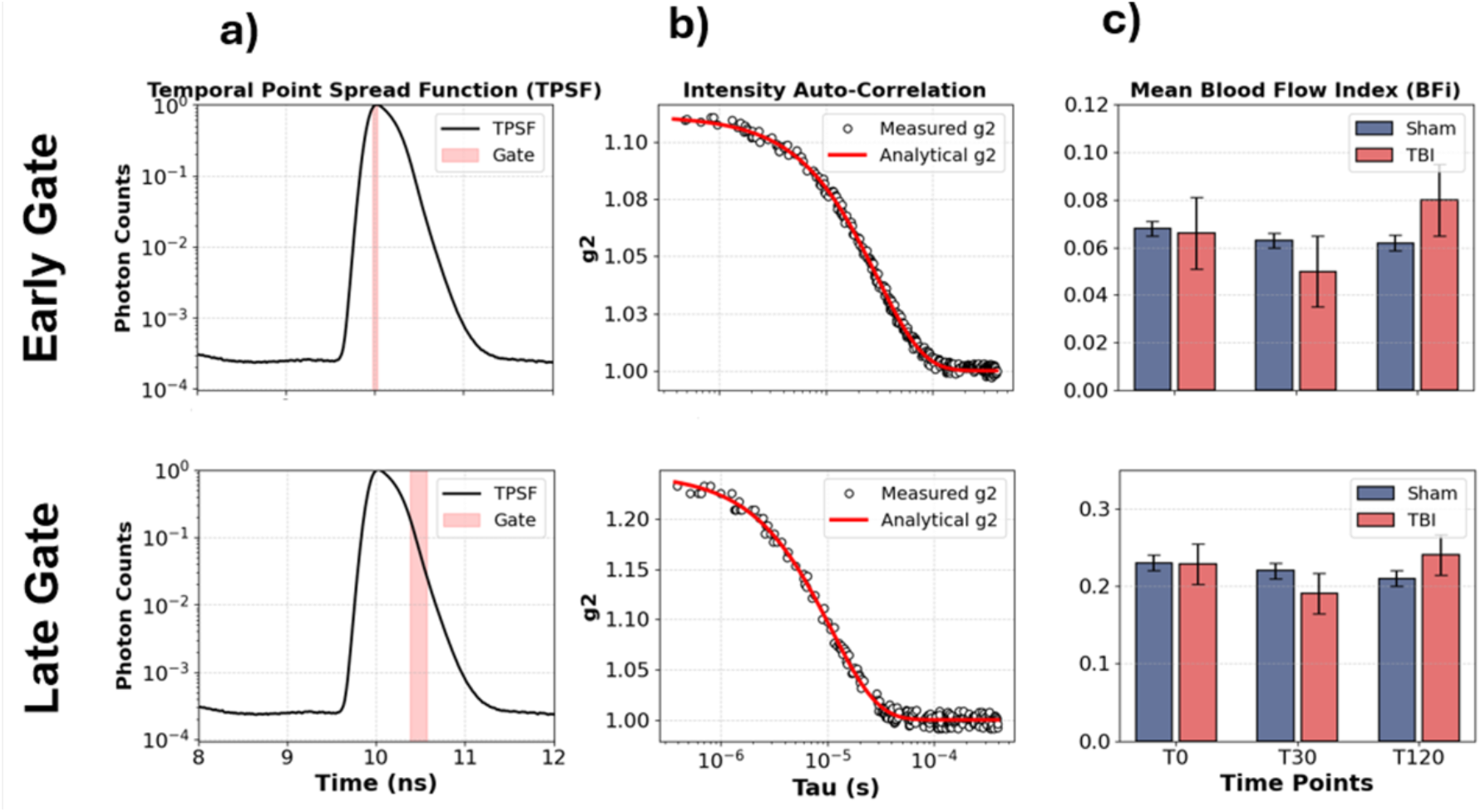
Representative results for temporal point spread function (TPSF) curves and mean blood flow index by TG-DCS in early and late gate. (a) This column shows TPSF curves in one subject with the early and late gate (EG, LG) highlighted. The EG opens -100ps with a width of 20ps from the peak and the LG opens -350ps with a width of 200ps from the peak (red overlay). (b) represents g2 curves of the subject that is fitted with the time-gated g2 model in early and late. (c) Measurement of BFI changes pooled across all mice in sham and injury groups starting before injury (T0) and continuing for half an hour (T30) and two hours (T120) in the early and late gates. BFI dropped within minutes of TBI, remained suppressed for 2 hours and gradually increased after 2 hours.

As the bar plot and Table-2 summarizes, for the TBI group BFI at the late gate was lower at T30 than baseline and gradually increased at T120 reaching baseline levels. More specifically, for the late-gate, for the TBI group, the baseline (T0) value ∼0.22±0.03, decreased to ∼0.19±0.02 at T30 and ∼0.24±0.03 at T120, and for the early-gate these numbers were 0.06±0.02, ∼0.05±0.02 and ∼0.08±0.02, respectively. Although there was a trend, these changes did not show statistical significance (as Table-3 summarizes), possibly due to limited number of mice.

**Table 1.** Frequency bands for low-frequency oscillations with respect to band-1 to band-5.

| Band | Frequency Range | Possible Mechanism |
| --- | --- | --- |
| V | 0.01-0.027 Hz | Endothelial (CA) |
| IV | 0.027-0.073 Hz | Neurogenic (activation/Brain function) |
| III | 0.073-0.198 Hz | Respiratory/ Brain function (lower ends) |
| II | 0.198-0.5 Hz | Mayer wavers (ABP)/Respiratory |
| I | 0.5-0.75 Hz | Mayer waves (ABP) |

**Table 2.** BFI values for traumatic brain injury (TBI) group vs. Sham group at three time points, T0: baseline, T30: 30-min post-trauma, and T120: 120-min post-trauma.

| Time Point | Early Gate |  | Late Gate |  |
| --- | --- | --- | --- | --- |
|  | Sham | TBI | Sham | TBI |
| T0 | $0.07 \pm 0.01$ | $0.06 \pm 0.01$ | $0.23 \pm 0.04$ | $0.22 \pm 0.03$ |
| T30 | $0.06 \pm 0.01$ | $0.05 \pm 0.01$ | $0.22 \pm 0.03$ | $0.19 \pm 0.02$ |
| T120 | $0.06 \pm 0.01$ | $0.08 \pm 0.01$ | $0.21 \pm 0.02$ | $0.24 \pm 0.03$ |

**Table 3.**
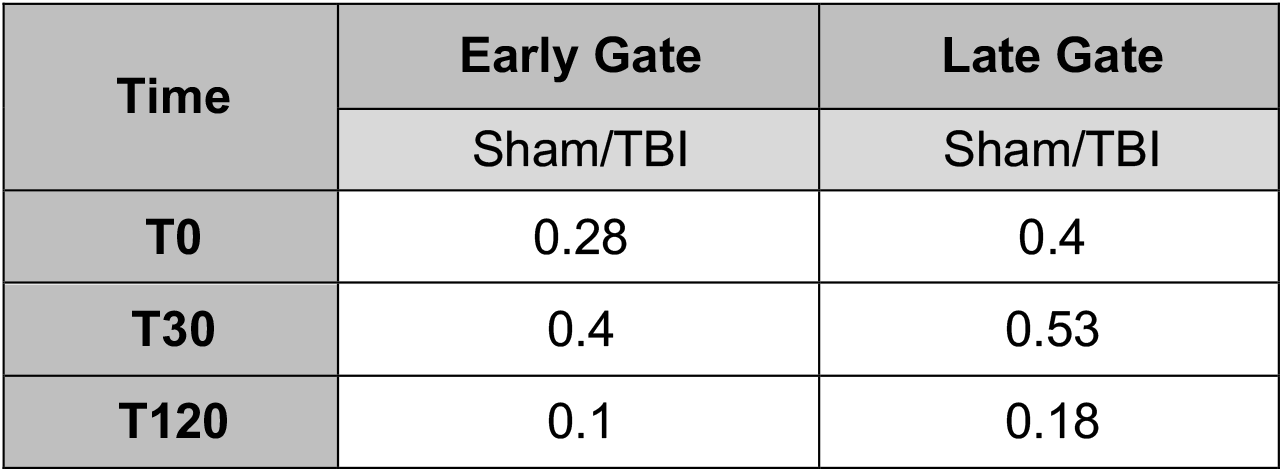
Statistical results (p-values) for the changes in BFI.

### 2.2. Frequency-domain characteristics

Next, we evaluated the LFOs in tissue blood flow measured by TG-DCS. For this purpose, we calculated the meantime course of BFI across all TBI data at 3 time points after detrending and normalizing for both gates as shown in Figures 3 (a) and (c). Then, we analyzed the power spectral density (PSD) changes with a passband at the LFO range from 0.01 to 0.15 Hz by comparing the PSD changes of different frequency bands, following baseline, post-30 minutes and post-120 minutes time points on both gates (Fig 3 (b) and (d)).

**Figure 3.**
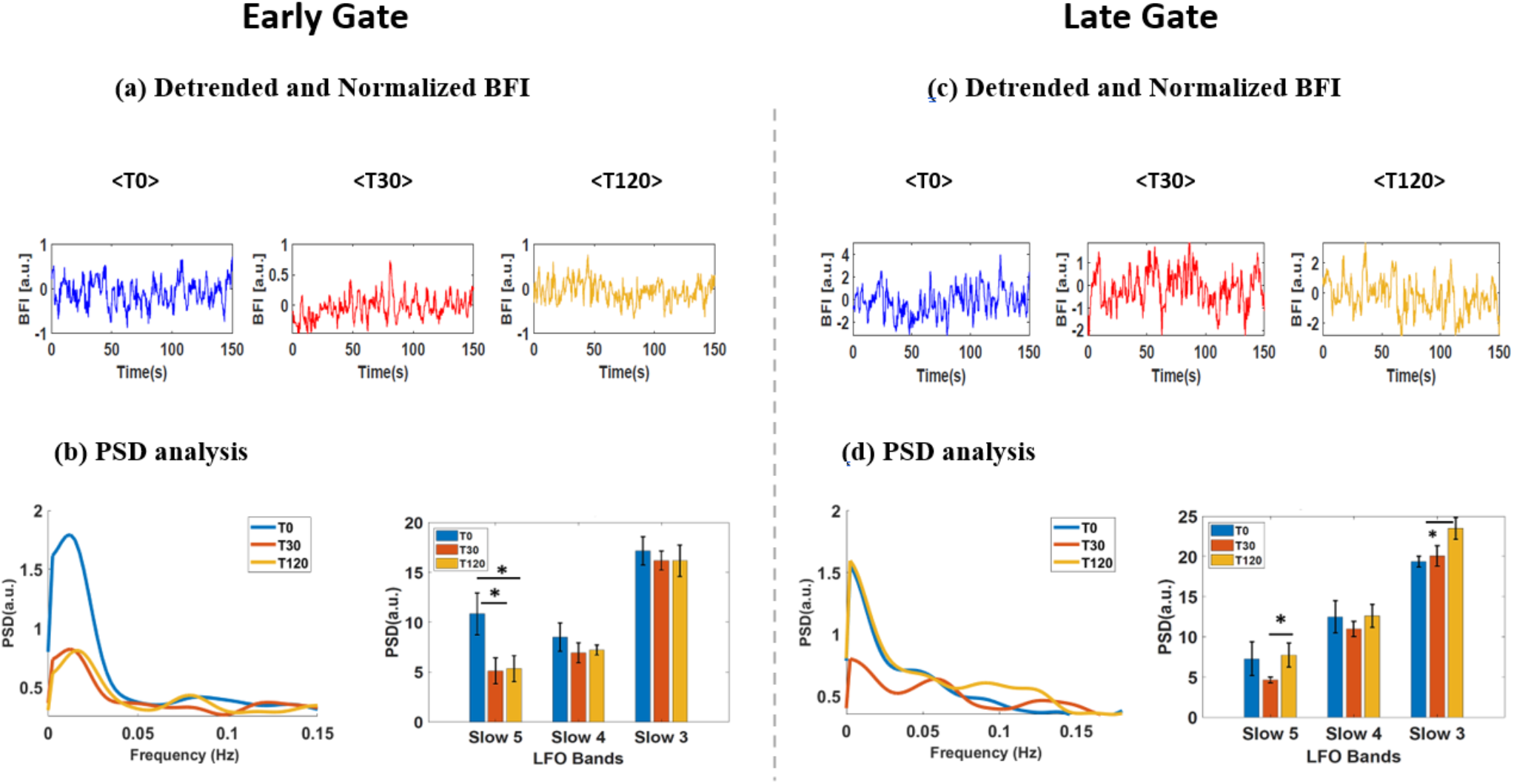
Representative of BFI changes and PSD analysis of Low-Frequency Oscillation (LFO). (a and c) Mean time course of BFI across all TBI data, according to the T0, T30, and T120 periods in early and late gate. (b and d) Relative PSD analysis among three frequency bands based on the T0, T30, and T120 periods. Detrended and normalized PSD distributions among 3 frequency bands based on baseline (T0: blue), after 30 minutes (T30: red), and 2 hours after impact (T120: orange) timepoints across all TBI data in the early and late gate. The p-values were compared by rank sum across subjects (paired one-sided, p< 0.05 indicated by * in both gates). The PSD of EG showed significant changes in slow band-5 between T0/T30 and T0/T120. The LG showed significant changes in bands 5 and 3 between T30/T120.

As shown in Fig 3 (b) for the early gate and for the baseline period (T0), the relative PSDs for Slow-5, -4 and -3 bands were 10.84, 8.51, and 17.16 [a.u.], respectively. For the relative PSDs of bands 5, 4, and 3 in T30, the values were 5.11, 6.93, and 16.18 [a.u.], respectively. In the T120 period, the relative PSDs were 5.34, 7.22, and 16.16 [a.u.], respectively, showing recovery to the baseline. In the T30 and T120 periods, the relative PSDs of band 5 substantially decreased and then increased after 2 hours, but at a lower level of the baseline. Fig 3 (d) for late gate, for the baseline period (T0), the relative PSDs for bands 5, 4, and 3 were 7.27, 12.49, and 19.34 [a.u], respectively. For the relative PSDs of bands 5, 4, and 3 in T30, the values were 4.63, 10.97, and 20.06 [a.u.], respectively. In the T120 period, the relative PSDs for the LFOs were 7.73, 12.62, and 23.5 [a.u.], respectively. In the T30 and T120 periods, the relative PSDs of bands 5 and 3 significantly decreased and increased after 120 minutes and slightly more than the baseline level. The significant p-value is indicated by * for both gates.

As **Table 4** indicates, we observed that the quantifying BFI at discrete time points in both gates showed significant changes in TBI relative to sham animals. For the early-gate, the rank sum test showed a significance with p-value= 0.01 for the changes between T0 and T30 and for the changes between T0 and T120 in LFO-V (band-5). For the late-gate, the rank sum test demonstrated significant changes with p-values of 0.03 and 0.04 for T30 and T120 in LFO-V and III (bands 5 and 3). Overall, as Figs 3(b) and 3(d) and Table 4 indicate, the TG-DCS system measures vasomotion tone within tissues in the VLF band to capture subtle changes in blood flow.

**Table 4.**
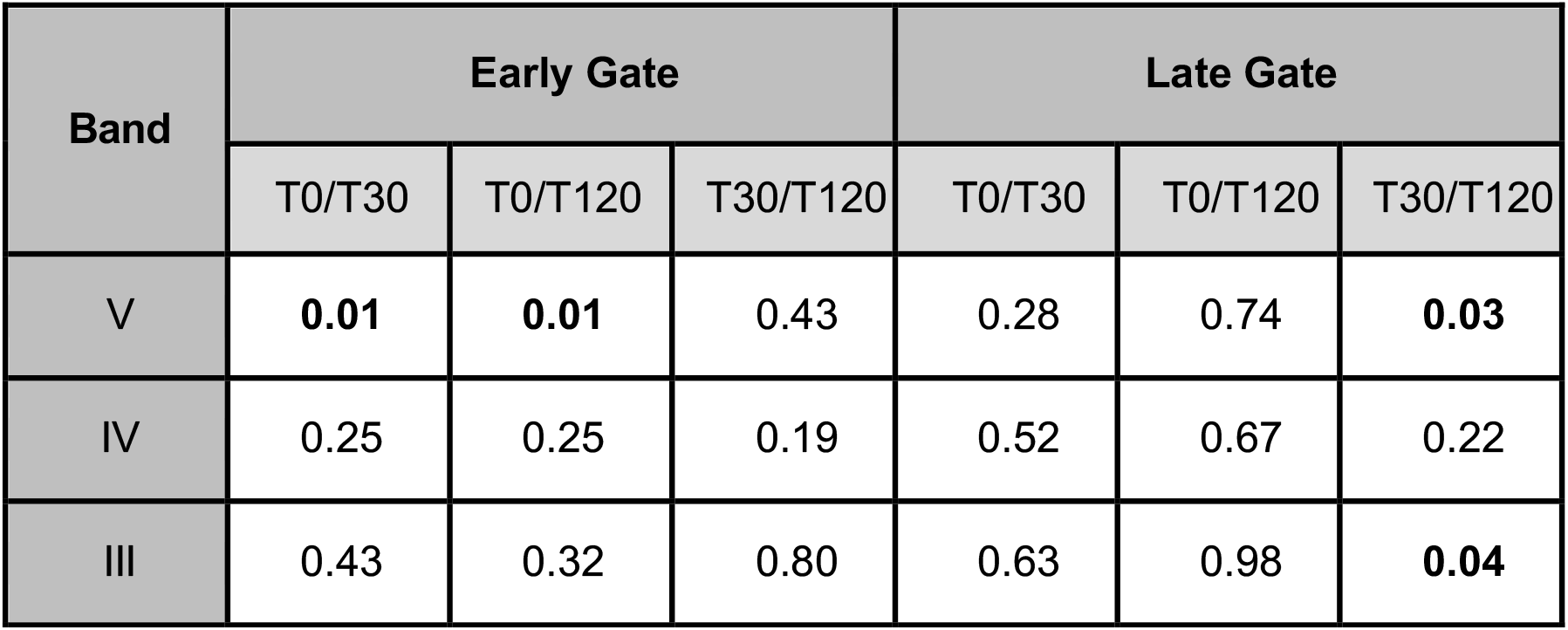
Statistical results for Low-Frequency Oscillations (LFOs). The PSD of EG shows significant changes in slow band 5 between T0/T30 and T0/T120 time-windows and for the LG through bands 5 and 3 between the T30/T120 time-windows.

| Band | Early Gate |  |  | Late Gate |  |  |
| --- | --- | --- | --- | --- | --- | --- |
|  | T0/T30 | T0/T120 | T30/T120 | T0/T30 | T0/T120 | T30/T120 |
| V | **0.01** | **0.01** | 0.43 | 0.28 | 0.74 | **0.03** |
| IV | 0.25 | 0.25 | 0.19 | 0.52 | 0.67 | 0.22 |
| III | 0.43 | 0.32 | 0.80 | 0.63 | 0.98 | **0.04** |

## 3. Discussion

Time-gated diffuse correlation spectroscopy (TG-DCS) offers a powerful, non-invasive method for measuring cerebral blood flow (CBF) and its oscillatory features at different tissue depths. By leveraging time-of-flight photon separation, TG-DCS enables depth-sensitive assessment of cerebral hemodynamics, distinguishing superficial (scalp) and deeper (cortical) signals. However, the ability of DCS is hindered by the low signal-to-noise ratio (SNR) when measuring blood flow in deeper tissue, since the fraction of detected photons decreases significantly with depth^10,13,49– 51^. Others and we showed that SNR can be improved improvement by performing measurements at 1064 nm by using superconducting nanowire single photon detector (SNSPD) detectors ^17–19,21,52^ . In our previous work, we demonstrated blood flow measurements of traumatic brain injury patient with 50 Hz temporal resolution in a clinical setting^18^. In this work, we applied this approach to monitor the depth-sensitive blood flow changes in mice after injury challenge. Because of smaller skull thickness (<1 mm vs. ∼10mm in humans) and shorter source-detector separation (4 mm vs. 15 mm for humans) used, the SNR was sufficient to implement narrow time-gating to separate superficial vs. deeper signals, aligns well with previous work by Sutin *et al*. ^13^ and by Samaei *et al*.^53^ Although blood flow at early and late gates decreased, these changes did not show statistical significance, potentially due to the small number of animals. Each animal may have potentially higher impact from the surface due to drop off the ball and may induce global changes. These results align well with the previous studies by Abookasis *et al*. and Witkowski *et al*., who also observed a similar decreasing trend by using the superficial optical speckle imaging method ^54,55^. We reported a significant (∼15%) decrease within 30 min of continuous measurements by using the speckle contrast optical spectroscopy (SCOS) method ^56^, which is originally based on the DASCA method developed by the Lee group ^41,57–59^. Jang *et al*. observed an acute blood flow decrease using DCS within 30 min ^60^, while Buckley *et al*. also used DCS for monitoring repetitive concussions in mice for 24 h and monitored several days and observed an initial cerebral blood flow (CBF) decrease at 4 h ^61^.

Beyond mean blood flow, we examined low-frequency oscillations (LFOs) in the BFI signal, as these have been associated with autoregulation, vasomotion, and neurovascular coupling ^33,60,62– 64^. Power spectral density (PSD) analysis revealed significant alterations in LFO bands after injury, particularly in slow-5 (0.01–0.027 Hz) and slow-3 (0.073–0.198 Hz), at both gates. These findings suggest that LFOs may be more sensitive than BFI alone in capturing the vascular consequences of TBI. Altered LFO patterns likely reflect disruptions in endothelial function, autoregulatory capacity, and vascular tone, all key components of the secondary injury cascade. Importantly, the differential LFO response in superficial versus deeper tissue underscores the value of depth-sensitive analysis.

Taken together, our results support the use of TG-DCS as a dual-domain tool: it enables both traditional blood flow quantification and frequency-domain analysis of physiological oscillations, which together may offer complementary insights into injury severity and progression. These findings align with prior reports that LFO metrics may serve as early indicators of cerebral dysfunction in TBI and other neurovascular disorders. These observations are consistent with other research indicating that TBI induces profound and sustained disruptions in blood flow, which are critical to the pathology of brain injuries. Furthermore, the differential response in blood flow and its oscillations between superficial and deeper cortical layers underscores the importance of depth-sensitive techniques in TBI research.

Our results complement and extend our prior clinical demonstration of TG-DCS in the neurocritical care setting (Poon et al.^65^), where we achieved bedside measurements of cerebral blood flow using a similar 1064 nm system with SNSPD detectors. In the present preclinical study, we adapted the same platform to investigate early physiological responses to TBI in a controlled animal model, enabling finer control of measurement conditions and trauma timing. Importantly, we demonstrate that TG-DCS not only quantifies depth-resolved blood flow but also captures low-frequency oscillations (LFOs) that appear sensitive to neurovascular disruption. These findings establish the utility of TG-DCS in characterizing early autoregulatory dysfunction and lay the groundwork for its use as a translational physiological biomarker platform in both animal and human TBI studies.

Despite its strengths, the current study has several limitations. The use of a mouse model, while widely used, may not fully capture the complexities and heterogeneity of traumatic brain injury observed in humans. Differences in cerebral anatomy, physiology, and injury mechanisms can affect the translatability of findings. The mouse head, while a complex structure, treated here as a homogeneous medium. The experimental results thus reflect average values of selected gates related to superficial vs. deeper regions. Time-gated diffuse correlation spectroscopy has limitations in distinguishing exact contributions from different microvascular compartments within the brain compared to high-resolution imaging modalities like microscopy and optical coherence tomography (e.g., ^66–70^), and mesoscopy (e.g., ^71–74^). Continuous monitoring could also prove beneficial for analyzing post-impact kinetics rather than three discrete time points ^55,56,60^ . Further, our study’s duration was confined to 120 minutes post-trauma to assess the early changes, whereas extending to longer periods could mimic broader clinical conditions ^61^. Our optical method can provide longitudinal monitoring over extended hours and days; thus, this aim can be pursued in future studies. Moreover, since the signal is averaged across a volume, adopting an imaging approach could enhance the assessment of specific traumatic changes at local positions, such as impact zone vs further distance areas ^55^. Although we did not attempt to quantify optical parameters, in principle they can be extracted by using instrument response function deconvolution methods^75^ . While we did not record systemic arterial blood pressure (ABP) concurrently, previous studies have demonstrated that spontaneous LFOs in cerebral hemodynamics reflect a combination of endothelial, neurogenic, and metabolic influences ^38,76^. Our depth-resolved LFO analysis, therefore, offers insight into potential disruptions in neurovascular regulation after TBI, even in the absence of explicit pressure-flow coupling metrics. Future studies would benefit from integrating systemic physiological monitoring and exploring methods to enhance penetration depth and resolution to obtain a more comprehensive view of the TBI impact across the entire brain.

## 4. Conclusions

In this study, we demonstrated the utility of time-gated diffuse correlation spectroscopy (TG-DCS) with SNSPD detection at 1064 nm for non-invasive, depth-sensitive assessment of cerebral blood flow (CBF) and low-frequency oscillations (LFOs) in a mouse model of traumatic brain injury (TBI). By separating superficial and deeper photon arrival times, we successfully monitored cortical hemodynamic responses to injury over time. Our findings revealed a decrease in blood flow within 30 minutes post-injury, followed by a partial recovery by 120 minutes, across both early and late gates. More notably, spectral analysis of BFI signals showed significant changes in LFO power, particularly in slow-5 and slow-3 frequency bands, indicating disruptions in vascular autoregulation and neurovascular coupling. These LFO alterations were more sensitive than BFI alone, suggesting their value as potential biomarkers for early neurovascular dysfunction after TBI. Together, this work highlights TG-DCS as a promising tool for depth-resolved, dynamic monitoring of cerebral perfusion and vascular oscillations, and lays the foundation for future preclinical and translational studies aimed at improving diagnosis and therapeutic monitoring in acute brain injury.

## 5. Materials and Methods

### 5.1 Animal Preparations and Experimental Protocols

All animal experiments followed protocols approved by the Wright State Department of Laboratory Animal Resources (LAR) and the Institutional Animal Care and Use Committee. All animal care was performed in accordance with the relevant guidelines and regulations outlined in the “Guide for Care and Use of Laboratory Animals”, and in accordance with ARRIVE guidelines. The weight drop technique was chosen to induce traumatic brain injury (TBI) as it closely mimics focal head trauma, and the optical setup was used to record changes in blood flow to study the effects of TBI on cerebral blood flow. The experiment involved a total of 12 female C57BL/6J mice (Jackson Lab, ME), which were divided into two groups: 4 control mice and 8 mice designated for the induction of TBI. Mice were anesthetized using 5% isoflurane in an anesthesia chamber for 180 seconds. After the initial anesthesia, mice were given 2.5% isoflurane and fitted with a nose cone to maintain sedation under anesthesia for a subsequent 180 seconds before starting the measurements. Baseline TG-DCS measurements were taken for the next 10 minutes before removing the nose cone and placing the mouse on the weight drop apparatus. In the experimental protocol, three periods were defined as baseline (T_-10_: 10 minutes before impact), T_30_ (30 minutes after impact), and T_120_ (120 minutes after impact).

The first step of the experiment involved taking control measurements in the control group of mice. These measurements were taken for a period of 10 minutes to establish a baseline measurement of blood flow. The weight drop protocol ^77–79^ was used to cause the TBI, which involved dropping a cylindrical metallic weight weighing 100g from a height of 90 cm on the head of the mouse, between the anterior coronal suture and the posterior coronal suture. After inducing TBI, the time taken for the mouse to wake up was measured before transferring it back to its cage. It was then taken out of the cage and anesthetized for 5min before the T30 timepoint. The optical measurements commenced for a duration of 10 minutes. The same procedure was followed at T120.

### 5.2 TG-DCS System for Data Acquisition

A portable prototype of TG-DCS was developed for clinical translation, as we detailed previously^18^. Here, we adapted the probe for preclinical studies. The probe consisted of a 1 mm multi-mode source fiber secured together with a single-mode fiber separated by 4 mm. The few-mode fiber was connected to the superconducting nanowire single photon detector (SNSPD; QOpus, Plymouth, MI) to collect reflected photons at 1064 nm, and was connected to a time-correlated single-photon counter (TCSPC) (HydraHarp400; Picoquant Inc, Berlin, Germany). The laser source consists of a 1064nm seed laser, operating at a repetition rate of 80MHz with a pulse width of 250ps (QLD-106P; QDLaser Inc, Kanagawa, Japan). The seed laser was amplified with a 1064nm optical amplifier (Mantaray-Amp-1064; Cybel LLC, Bethlehem, PA) fiber-coupled to the source fiber with an average power of ∼10mW. In time-domain DCS, the light from a pulsed laser is injected into the sample through the single-mode fiber, and the reflected light is detected with a SNSPD (**Fig.1**) ^13,16,18,19,49,51,53,80–82^ . The instrument response function (IRF) was measured by placing a thin scattering layer (Teflon Tape; 3M, Saint Paul, MN) between the source fiber and the detector fiber. All experimental measurements were performed in reflection geometry at source-detector separation, ρ = 4 mm. To ensure consistent conditions for all experiments and to minimize the environmental noise, measurements were performed in a dark room at a temperature of 25 °C.

### 5.3 Data Processing for Blood Flow

To measure TG-DCS, we utilized two tagged time points to estimate time gate selection and to determine the intensity autocorrelation function (ACF). The first tag is the micro-time, which corresponds to the photon time-of-flight (TOF). It is the time needed for the photon to travel from the source to the detector and is used to estimate the TPSF (temporal point spread function), which is then used to select a time-gate for photon measurement. The second tag is the macro-time, which corresponds to the absolute arrival time, or the time since the measurement was started. It is used to determine the intensity autocorrelation function (ACF). ACF is commonly defined as ^13,49,50^ :

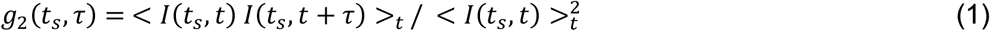

where t_s_ denotes the time-of-flight, τ is the autocorrelation delay/lag time, and < … >_t_ denotes temporal averaging with respect to t distinct from t_s_.

### Estimating diffusion coefficient and blood flow index

In most TG-DCS studies, the measured time-resolved intensity autocorrelation function, *g*_*2*_(*t*_*s*_, *τ*) is fit to the following model (Siegert relationship) ^13,49,50^ :

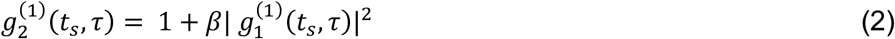

where g_1_^(1)^ (t_s_, τ) = exp [−ξ(t_s_)τ] is the single-exponential optical field autocorrelation function of the specific pathlength. The symbol ξ(t_s_) stands for the TOF-dependent ACF decay and the β term ranges from 0 to 1 for the spatial and temporal coherence of light in the experimental setup ^83,84^. Let’s first assume that we know the length of the path a photon traveled, s. The field autocorrelation function can then be calculated as ^13,49,50^ :

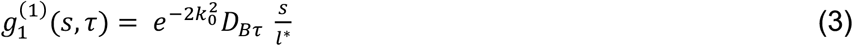

where 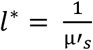 is the transport mean free path (i.e., the expected distance a photon travels before beginning scattered, analogous to the definition of 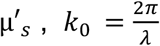 is the wavenumber of the light in the medium. That is, given a light of a certain wavelength and scattering angle impinging on a medium with a Brownian diffusion coefficient D_B_ and it is referred to as the blood flow index (BFi). The total field autocorrelation function is then a weighted average over all possible pathlengths, given by:

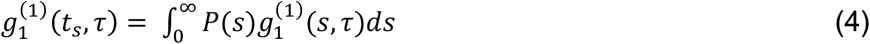

where P(s) is the temporal point spread function (TPSF) of the tissue in a specific gate, normalized such that it is a valid probability distribution, and where the pathlength can be simply related to the time-of-flight as s = t * v, where v is the speed of light in the medium. P(s) and therefore g_1_ will depend on the optical properties of the medium, where mice optical properties were estimated relative to the known phantom and used to fit the 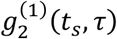 Eq.(2) to the experimentally estimated *g*_2_(*t*_*s*_, *τ*)Eq.(1). Also, we could limit our measurements to photons of known pathlength by time-gating to find P(s) by directly measuring the photon time-of-flight and use the pathlength-dependent autocorrelation function 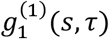 to estimate the dynamical properties, also known as the diffusion coefficient (D_B_). In this study, two gates were selected: the early gate (EG) (-100ps relative to the peak of each TPSF with a width of 100ps) and the late gate (LG) (+350ps relative to the peak of each TPSF with a width of 200ps). The EG and LG data were autocorrelated from 5e-7s to 1e-3s with an integration time of 0.1s.

#### General expression for the ACF

The measured TOF-resolved intensity autocorrelation function 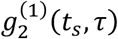 Eq. (2) depends on the instrument response function or the IRF and the photon time-of-flight distribution, when experimental estimates are integrated over the TOF. Thus, under the validity of Eq. (2) and Eq. (4), we can include those effects as follows: ^51,82,85^

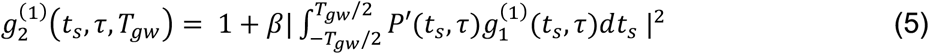

where *T*_*gw*_ is the gate width, and *P*^*′*^(*t*_*s*_) is the normalized measured photon distribution of time-of-flight (DTOF) via a convolution (*) with the IRF, *I*_0_(*t*_*s*_):

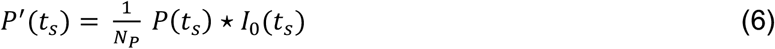

where 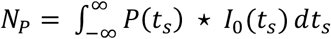 is the normalization factor.

Therefore, to improve the signal-to-noise ratio, the experimental estimates are integrated over the photon TOF range. Here, we only consider relatively narrow time gates, neglecting the IRF and TOF integration effects. For all reported data in this paper, *g*_*2*_(*t*_*s*_, *τ*) was obtained from a rectangular time gate with 20ps width for the early gate and 200ps width for the late gate.

### 5.4 Power Spectral Density Analysis for Low-Frequency Oscillations (LFOs)

In addition to the blood flow changes, we also investigated the power spectral density (PSD) analysis of the blood flow index to quantify the changes in low-frequency oscillations (LFOs). since LFOs can serve as a potentially novel metric for brain function, and optical blood flow can detect these oscillations ^23,25,42,56,60,86^. The frequency distribution of spontaneous LFOs are characterized by the following bands, labelled slow-5 (0.01–0.027 Hz), slow-4 (0.027–0.073 Hz), and slow-3 (0.073–0.198 Hz)^47,87–89^, where slow-5 can be endothelial or CA, while slow-4 neurogenic or brain activation, and slow-3 can be related to respiratory and brain function (lower ends like ∼0.1 Hz) mechanisms ^47,89^, as summarized in **Table-1**. Here, we mainly focused on the slow bads having lower frequencies (bands V, IV, and III), and presented the results related to these bands, since Bands I and II reflect non-cortical physiology.

As a more relevant to our optical approach, the power spectral density (PSD) analysis for LFOs intensity quantification has been previously implemented in diffuse optical technologies, including NIRS ^30,35,37,46,64,90–100^ and DCS ^34,45,101,102^. Following these approaches, LFO intensities of BFi at different time-points (i.e., baseline, 30 min after impact, and 120 min after impact) were extracted from PSDs by using Welch’s method in a nonparametric approach ^103^. The PSD was calculated as:

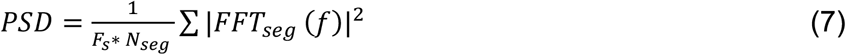

where Fs is the sampling frequency, N_seg_ is the number of segments, and FFT_seg_ is the fast Fourier transform of the segmented signals ^103^. Briefly, the time course of the BFI data under each physiological condition (T0, T30, T120) at the sampling rate of ∼10 Hz was first normalized and then detrended using a 2^nd^ order built-in “detrend” function in MATLAB (MathWorks Inc) to obtain the best fit for the trend. The detrended BFI data was then transferred into a 2^nd^ order zero-phase Butterworth filter with a passband at the LFO range from 0.01 to 0.15 Hz. Finally, the built-in “pwelch” function in MATLAB was used to generate the PSD over the LFO bandwidth. The PSD calculated across all subjects with (512 data points and 50% overlap). Then, the data was filtered into distinct frequency sub-bands to study frequency-specific CBF response.

## Data Availability

The datasets used and/or analysed during the current study are available from the corresponding author on reasonable request.

## Funding

NIH R01 (NIBIB Brain Initiative, 7R01EB031759-03).

## Acknowledgments

The authors acknowledge the funding support from NIH R01 (NIBIB Brain Initiative (7R01EB031759-03). We thank Ben Rinehart for the help with the animal handling during the measurements. We also thank Nick Bertone (Picoquant Inc.) for providing the demo unit of HydraHarp400 for the time domain acquisition system.

## Contributions

U.S. and B.F. conceived the experiments; C-S.P., T.M.R., and A.J.M. contributed to the preparation of the experimental system; C-S.P. and D.S.L. conducted the experiments, S.S. performed the data analysis; U.S., B.F. interpreted the results; S.S., U.S., B.F. and C-S.P. drafted the manuscript; all authors reviewed the manuscript.

## Competing interests

The authors declare no competing interests.

## Notes

### Competing Interest Statement

The authors have declared no competing interest.

